# Serum Neutralizing Activity of mRNA-1273 against SARS-CoV-2 Variants

**DOI:** 10.1101/2021.06.28.449914

**Authors:** Angela Choi, Matthew Koch, Kai Wu, Groves Dixon, Judy Oestreicher, Holly Legault, Guillaume B.E. Stewart□Jones, Tonya Colpitts, Rolando Pajon, Hamilton Bennett, Andrea Carfi, Darin K. Edwards

**Affiliations:** Moderna Inc., 200 Technology Square, Cambridge, Massachusetts, 02139, USA

## Abstract

The emergence of severe acute respiratory syndrome coronavirus 2 (SARS-CoV-2) variants has led to growing concerns over increased transmissibility and the ability of some variants to partially escape immunity. Sera from participants immunized on a prime-boost schedule with the mRNA-1273 COVID-19 vaccine were tested for neutralizing activity against several SARS-CoV-2 variants, including variants of concern (VOCs) and variants of interest (VOIs), compared to neutralization of the wild-type SARS-CoV-2 virus (designated as D614G). Results showed minimal effects on neutralization titers against the B.1.1.7 (Alpha) variant (1.2-fold reduction compared with D614G); other VOCs such as B.1.351 (Beta, including B.1.351-v1, B.1.351-v2, and B.1.351-v3), B.1.617.2 (Delta), and P.1 (Gamma) showed decreased neutralization titers ranging from 2.1-fold to 8.4-fold reductions compared with D614G, although all remained susceptible to mRNA-1273–elicited serum neutralization.

## INTRODUCTION

As the coronavirus disease 2019 (COVID-19) pandemic continues to escalate in various parts of the world, several severe acute respiratory syndrome coronavirus 2 (SARS-CoV-2) variants of interest (VOIs) and variants of concern (VOCs) have emerged, including in the United States (B.1.526; Iota), United Kingdom (B.1.1.7; Alpha), Brazil (P.1; Gamma), India (B.1.617.1, Kappa; B.1617.2, Delta), South Africa (B.1.351; Beta), Uganda (A.23.1), Nigeria (B.1.525; Eta), and Angola (A.VOI.V2).^1^ There is growing concern over these variants based on increased transmissibility and the ability of some variants to partially escape both natural and vaccine-induced immunity. Notably, the B.1.617.2 lineage was recently classified as a VOC by the World Health Organization due to evidence of an increased rate of transmission, reduced effectiveness of monoclonal antibody treatment, and reduced susceptibility to neutralizing antibodies.^1^

We previously reported that mRNA-1273, a lipid nanoparticle encapsulated mRNA-based vaccine encoding the spike glycoprotein of the SARS-CoV-2 Wuhan-Hu-1 isolate, induced high neutralizing antibody titers in phase 1 trial participants^2^ and was highly effective in preventing symptomatic and severe COVID-19.^3,4^ Some VOCs or VOIs, including B.1.351 and P.1, reduced neutralizing antibody levels using a pseudovirus-based model.^5^ Importantly, however, all variants remained susceptible to mRNA-1273 vaccine–elicited serum neutralization.^5^ Here we provide an update on the neutralization activity of vaccine sera against several newly-emerged variants, including the Delta variant B.1.617.2.

## METHODS

### Clinical Trial

Participants were immunized with 100 µg mRNA-1273 on a prime-boost schedule and sera was collected 7 days post boost (day 36). Study protocols and results are reported previously.^2^

### Recombinant vesicular stomatitis virus (VSV)-based pseudovirus assay

Codon-optimized full-length spike (S) protein of the original Wuhan-Hu-1 variant with D614G mutation (D614G) or the S variants, listed in **Table 1**, were cloned into a pCAGGS vector. To make SARS-CoV-2 full-length S pseudotyped recombinant VSV-ΔG-firefly luciferase virus, BHK-21/WI-2 cells (Kerafast) were transfected with the S-expression plasmid and subsequently infected with VSVΔG-firefly-luciferase as previously described.^6^ For the neutralization assay, serially diluted serum samples were mixed with pseudovirus and incubated at 37 °C for 45 minutes. The virus-serum mix was subsequently used to infect A549-hACE2-TMPRSS2 cells^7^ for 18 hours at 37 °C before adding ONE-Glo reagent (Promega) for measurement of the luciferase signal by relative luminescence units (RLUs). The percentage of neutralization was calculated based on the RLUs of the virus-only control, and subsequently analyzed using four-parameter logistic curve in Prism v.8 (GraphPad Software, Inc.).

**Table 1.**
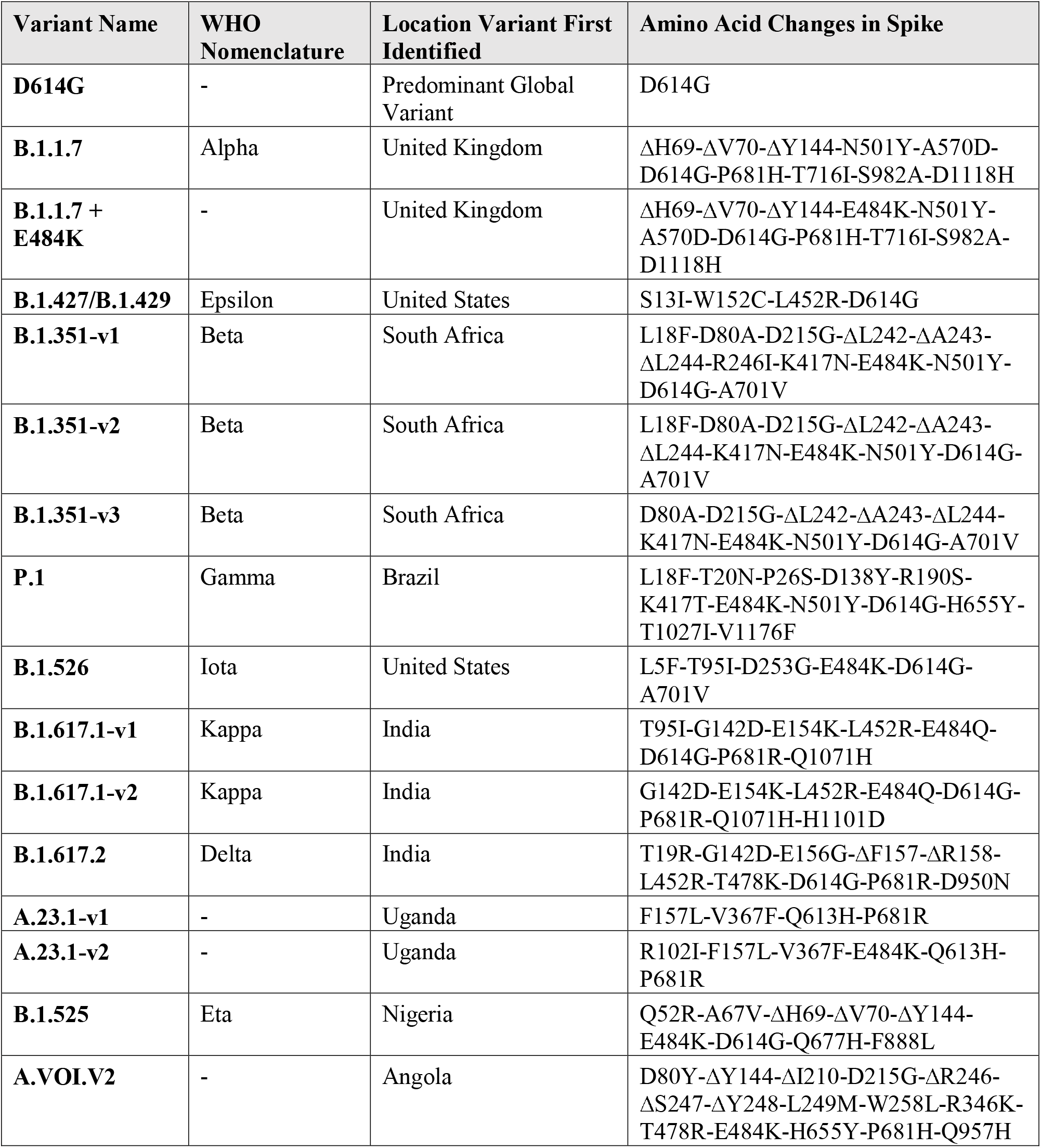
Spike mutations in SARS-CoV-2 variants evaluated in this study.

### Statistical Analysis

Two-sided Wilcoxon matched-pairs signed rank test was used to compare the same patients against different viruses. Statistical analyses were performed (Prism v.8). Geometric mean titers, lower limit of quantification, and fold change to D614G were included.

## RESULTS

We assessed neutralization activity of sera against D614G pseudovirus (predominant variant in 2020), B.1.1.7, B.1.1.7 + E484K, B.1.427/B.1.429, P.1, B.1.351-v1, B.1.351-v2, B.1.351-v3, B.1.526, B.1.617.1-v1, B.1.617.1-v2, B.1.617.2, A.23.1-v1, A.23.1-v2, B.1.525, and A.VOI.V2 (**Table 1**). Sera from the phase 1 mRNA-1273 clinical trial (8 participants, 1 week following Dose 2) were evaluated against each variant.^2^ Results showed minimal effects on neutralization titers against B.1.1.7 and A.23.1-v1 compared to D614G (**Figure 1**). In contrast, all other variants examined showed decreased neutralization titers compared with D614G (**Figure 1**), although all remained susceptible to mRNA-1273–elicited serum neutralization. Reductions in neutralization titers for these variants ranged from a factor of 2.1 to 8.4 compared with D614G (**Figure 1A**). Across the 3 versions of the B.1.351 variant tested, 6.9-fold to 8.4-fold reductions in neutralization were observed compared with D614G (**Figure 1A)**. Among all variants tested, the greatest effect on neutralization was observed for A.VOI.V2 and B.1.351-v3 (8.0-fold and 8.4-fold reductions compared with D614G, respectively). More modest 3.2- and 2.1-fold reductions were observed for P.1 and B.1.617.2, respectively. mRNA-1273 elicited neutralization titers against B.1.1.7, B.1.1.7+E484K, B.1.427/B.1.429, P.1, and B.1.351-v1 observed herein corroborated previous findings.^5^

**Figure 1.**
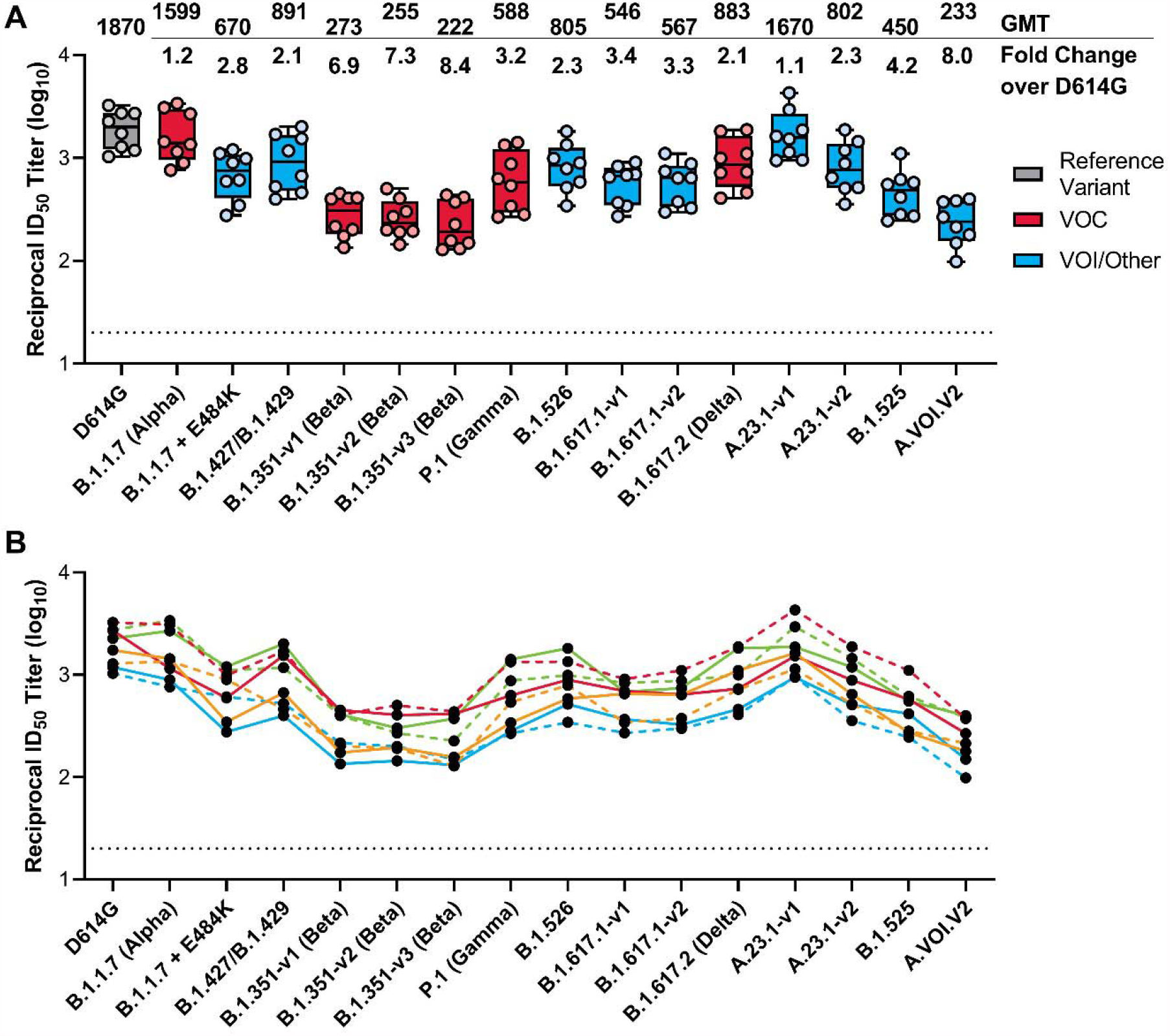
Neutralization of SARS-CoV-2 Pseudoviruses in Serum Samples. Serum samples were obtained from participants in the mRNA-1273 vaccine phase 1 trial on day 36 (7 days post dose 2). A recombinant vesicular stomatitis virus–based pseudovirus neutralization assay was used to measure neutralization. The pseudoviruses tested incorporated D614G or the spike substitutions present in B.1.1.7 (Alpha), B.1.1.7 + E484K, B.1.427/B.1.429, P.1 (Gamma), B.1.351-v1 (Beta), B.1.351-v2 (Beta), B.1.351-v3 (Beta), B.1.526, B.1.617.1-v1, B.1.617.1-v2, B.1.617.2 (Delta), A.23.1-v1, A.23.1-v2, B.1.525, and A.VOI.V2 (Table 1). The reciprocal neutralizing titers on the pseudovirus neutralization assay at a 50% inhibitory dilution (ID_50_) are shown. In Panel A, boxes and horizontal bars denote the interquartile range and the GMT, respectively. Whisker end points are equal to the maximum and minimum values below or above the median at 1.5 times the IQR. The GMT fold change over D614G for each variant is shown below. In Panel B, the colored lines connect the D614G and variant neutralization titers in matched samples. In both panels, the dots represent results from individual serum samples and the black horizontal dotted line represents the lower limit of quantification for titers at 20 ID_50_. GMT, geometric mean titer; ID_50_, 50% inhibitory dilution factor. VOC, variant of concern; VOI, variant of interest.

## CONCLUSION

Among VOCs tested, serum-elicited neutralization of the B.1.1.7 (Alpha) variant was comparable to D614G; a range of reduced neutralization titers compared to D614G were observed for other VOCs including the B.1.351 (Beta), B.1.617.2 (Delta), and P.1 (Gamma) variants, with reductions ranging from 2.1-fold to 8.4-fold. These data emphasize the need to continually assess the ability of mRNA-1273 to confer protection against prevalent and emergent VOIs/VOCs. Such data are crucial to inform necessary modifications to COVID-19 mRNA vaccines going forward, which may help to mitigate the ongoing spread of SARS-CoV-2 and the emergence of new variants.

## CONTRIBUTIONS

Conceptualization: K.W., A.C., M.K., J.O., G.S.J., H.L., R.P., A.C., and D.K.E.; methodology: K.W., A.C., M.K.; formal & statistical analysis: K.W., A.C., M.K.; writing—review and editing: K.W., M.K., A.C., G.D., G.S.J., T.C., H.B., A.C., and D.K.E. All authors have read and agreed to the published version of the manuscript.

## ACKNOWLEDGMENTS

This work used samples from the phase 1 mRNA-1273 study (NCT04283461; DOI: 10.1056/NEJMoa2022483). The mRNA-1273 phase 1 study was sponsored and primarily funded by the National Institute of Allergy and Infectious Diseases (NIAID), National Institutes of Health (NIH), Bethesda, MD; in part with federal funds from the NIAID under grant awards UM1AI148373, to Kaiser Washington; UM1AI148576, UM1AI148684, and NIH P51 OD011132, to Emory University; NIH AID AI149644, and contract award HHSN272201500002C, to Emmes. Funding for the manufacture of mRNA-1273 phase 1 material was provided by the Coalition for Epidemic Preparedness Innovation. We thank Dr. Michael Brunner and Dr. Michael Whitt for kind support on recombinant VSV-based SARS-CoV-2 pseudovirus production. Medical writing and editorial assistance were provided by Srividya Ramachandran, PhD, and Jared Cochran, PhD, of MEDiSTRAVA in accordance with Good Publication Practice (GPP3) guidelines, funded by Moderna Inc, and under the direction of the authors.

